# Single cell profiling to determine influence of wheeze and early-life viral infection on developmental programming of airway epithelium

**DOI:** 10.1101/2024.07.08.602506

**Authors:** Sergejs Berdnikovs, Dawn C Newcomb, Kaitlin E McKernan, Shelby N Kuehnle, Nana-Fatima Haruna, Tebeb Gebretsadik, Christopher McKennan, Siyuan Ma, Jacqueline-Yvonne Cephus, Christian Rosas-Salazar, Larry J Anderson, James E Gern, Tina Hartert

**Author notes:** co-first (Sergejs Berdnikovs, Dawn C Newcomb). contributed equally (Sergejs Berdnikovs, Dawn C Newcomb).

## Abstract

Although childhood asthma is in part an airway epithelial disorder, the development of the airway epithelium in asthma is not understood. We sought to characterize airway epithelial developmental phenotypes in those with and without recurrent wheeze and the impact of infant infection with respiratory syncytial virus (RSV). Nasal airway epithelial cells (NAECs) were collected at age 2-3 years from an *a priori* designed nested birth cohort of children from four mutually exclusive groups of wheezers/non-wheezers and RSV-infected/uninfected in the first year of life. NAECs were cultured in air-liquid interface differentiation conditions followed by a combined analysis of single cell RNA sequencing (scRNA-seq) and *in vitro* infection with respiratory syncytial virus (RSV). NAECs from children with a wheeze phenotype were characterized by abnormal differentiation and basal cell activation of developmental pathways, plasticity in precursor differentiation and a delayed onset of maturation. NAECs from children with wheeze also had increased diversity of currently known RSV receptors and blunted anti-viral immune responses to *in vitro* infection. The most dramatic changes in differentiation of cultured epithelium were observed in NAECs derived from children that had both wheeze and RSV in the first year of life. Together this suggests that airway epithelium in children with wheeze is developmentally reprogrammed and characterized by increased barrier permeability, decreased antiviral response, and increased RSV receptors, which may predispose to and amplify the effects of RSV infection in infancy and susceptibility to other asthma risk factors that interact with the airway mucosa.

**SUMMARY:** Nasal airway epithelial cells from children with wheeze are characterized by altered development and increased susceptibility to RSV infection.

## Introduction

Childhood asthma is in part an airway epithelial developmental disorder and its origin and clinical manifestations are tightly linked with altered airway epithelial cell (AEC) physical, metabolic, and functional barrier properties. The airway epithelium consists of multiple specialized cell subsets forming a functional barrier against the external environment. Many of the risk genes for asthma are expressed in the airway epithelium, particularly allergy and epithelial barrier function genes, which support the importance of the airway epithelium in asthma development (*1, 2*). AEC barriers in childhood are shaped and regulated by active on-going developmental programs, with morphogenesis of lung epithelial cells and barrier function continuing throughout normal postnatal development (*3–7*). AEC differentiation occurs largely in the first year of life but continues until approximately two years of age. Asthma results from host and environment interactions, and environmental asthma risk factors such as pollution, tobacco smoke and respiratory viruses, interact directly with the developing airway epithelium (*8*). Since AEC development continues after birth, early-life mucosal environmental exposures have the unique opportunity to alter the course of AEC differentiation and contribute to asthma development.

Respiratory syncytial virus (RSV) is an airway mucosal pathogen which directly infects the airway epithelium and is one of the most consistently identified asthma risk factors with a high population-attributable fraction (*9, 10*). Respiratory viruses represent a unique early life exposure as they integrate into the cell in order to replicate, in contrast to inhalant and irritant exposures. RSV serves as an ideal model to understand the impact of early life environmental exposures on airway epithelial development in both *in vivo* and *in vitro* studies, as about half of infants are infected with RSV in the first year of life, providing comparator groups of infected and uninfected infants (*10, 11*). Airway epithelial changes have been demonstrated in children before the onset of asthma, suggesting that airway epithelial developmental changes occur early in asthma pathogenesis. However, single cell profiling of wheeze, which typically precedes asthma, and the impact of early life mucosal respiratory viruses such as RSV on epithelial developmental phenotype have never been studied (*12*).

The objective of this study was to test whether: (1) there are unique developmental characteristics of the early life airway epithelium that characterize childhood wheeze, including that the wheeze developmental phenotype manifests in air-liquid interface differentiation culture conditions, and (2) early life asthma risk factors, using natural infant RSV infection as a model, affect AEC developmental programming and AEC differentiation in culture. As the collection of infant and child bronchial epithelium is highly invasive and impractical in population-based studies, we utilized the nasal airway epithelium to characterize developmental phenotypes in health and disease. The nasal transcriptome has been demonstrated to be an excellent and well-accepted proxy of expression changes in the lung airway transcriptome in asthma, as well as in distinguishing phenotypes of asthma (*13, 14, 15*). We used nasal airway epithelial cells cultured in air-liquid interface differentiation conditions, in combination with single cell RNA sequencing (scRNA-seq) and *in vitro* infection with RSV to investigate AEC developmental phenotypes at the age of 2-3 years that characterize children with wheeze, and to determine if an early life environmental asthma risk factor, RSV, is associated with altered nasal airway epithelial cell (NAEC) development.

## Results

### Selection of study population based on wheeze and RSV exposures in infancy

To determine if wheeze and early-life (before one year of age) RSV infection are associated with altered AEC development, we collected NAECs from children ages 2-3 years, which represents the end point of postnatal differentiation trajectory, in the INSPIRE birth cohort. INSPIRE is a population-based birth cohort of healthy, term and predominantly non-low birthweight infants. The sub-group for this study was selected from an *a priori* designed nested cohort of 100 participants using a random number generator from 4 pre-defined groups based on their history of wheezing and confirmed presence/absence of RSV infection during the first year of life.

NAECs at age 2-3 years were collected from these four *a priori* groups: control (no wheeze/no infant RSV infection), RSV in the first year of life (no wheeze/infant RSV infection), wheeze starting in the first year of life (wheeze/no infant RSV), and wheeze and RSV in the first year of life (wheeze/RSV) (**Fig. S1**). Wheeze was tracked annually using a validated questionnaire, and RSV infection by age one year was defined with a combination of passive and active surveillance with viral identification through molecular and serological testing to identify RSV infection (*16, 17*). Two to three participants from each of these four mutually exclusive groups were randomly selected, balanced by sex for single cell RNA-sequencing, and five to six different samples were selected from each group for in vitro experiments. **Fig. 1** illustrates the use of NAECs that were collected and cultured at air-liquid interface (ALI) to allow for differentiation from the four *a priori* defined groups and single cell analysis. In additional experiments, transepithelial electrical resistance (TEER) was measured during NAEC differentiation, and NAECs from the four *a priori* defined groups were infected with a clinically relevant strain of RSV, RSV 01-2/20, and mock to measure RSV infectivity.

**Fig. 1.**
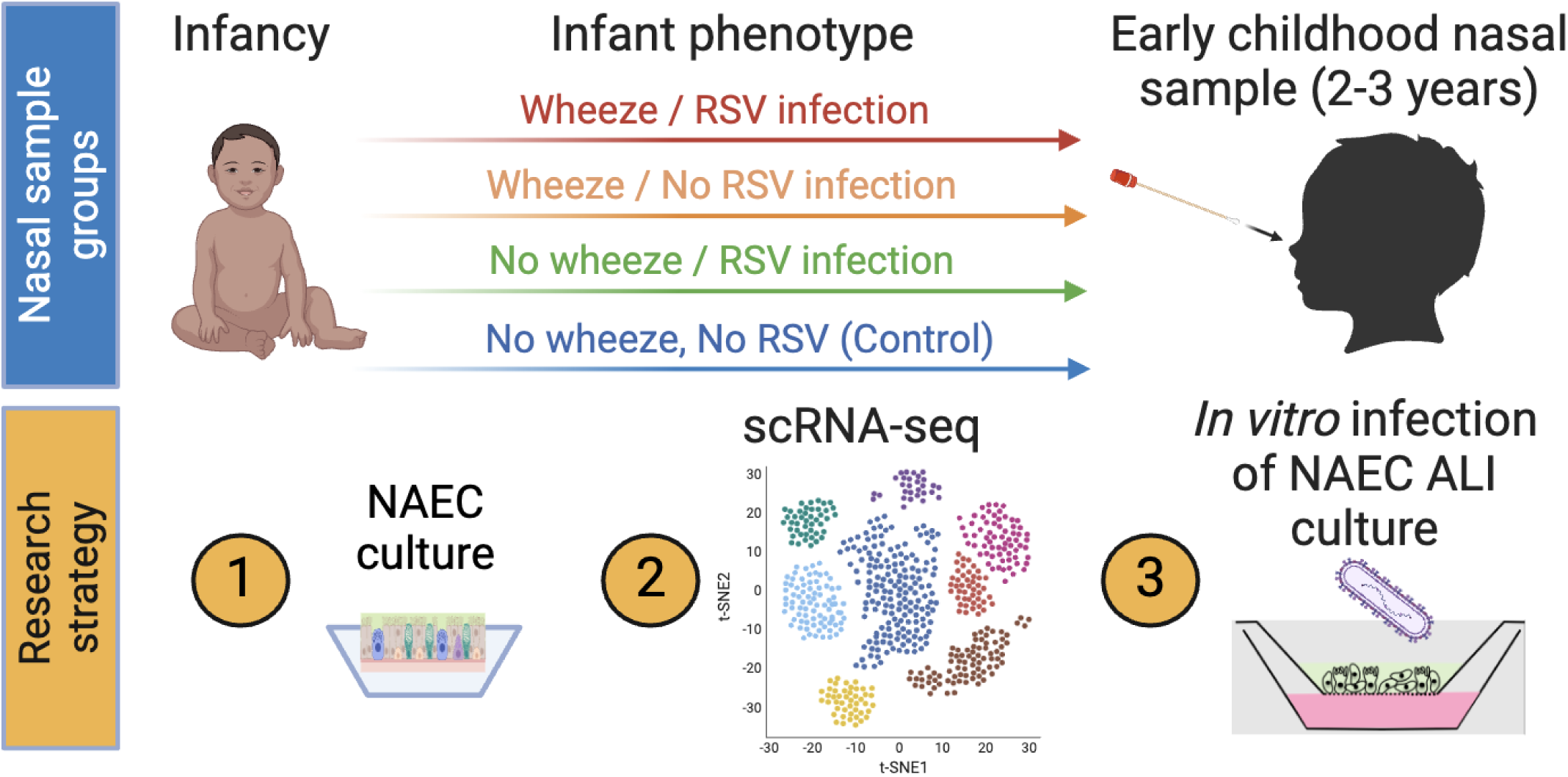
Graphical outline of research study workflow. (NAEC = nasal airway epithelial cell; scRNA-seq = single-cell RNA sequencing; ALI = air liquid interface)

### scRNA-seq analysis of differentiated NAECs from 2–3-year-olds identifies epithelial subset composition by wheeze and RSV phenotypes

First, we determined epithelial subset composition of the NAECs collected from children between age 2-3 years and cultured in ALI. We performed an integration scRNA-seq analysis of all epithelial samples and determined cell subset composition based on known markers of epithelial differentiation as published in Viera-Braga et al. (**Fig. 2B**) (*18*). We identified 17 different epithelial clusters across the four study groups, including basal, progenitor, and mucociliary subsets (**Fig. 2A**). Analysis of top marker genes differentially expressed across the seventeen clusters further confirmed functional identity of these subsets (**Fig. 2C**). Despite shared subset identity, several subsets of epithelial cells clustered separately based on group origin (i.e., three distinct clusters of club cells, two clusters of goblet cells and various precursors), likely driven by functional or developmental differences in the AEC phenotypes (**Fig. 2A and B**). Next, we compared subset composition of epithelial cells across the four pre-specified groups: (1) no wheeze/no RSV infection in infancy (control, -/-), (2) no wheeze/RSV infection in infancy (-/+), (3) wheeze/no RSV infection in infancy (+/-), and (4) wheeze/RSV infection in infancy (+/+). **Fig. 2D** shows cell integration from all four study group samples summarizing cell subset differences by group. There were appreciable differences in basal, precursor and club cells across different groups, especially in wheeze groups, while the RSV no wheeze group appeared more similar in epithelial composition to the control group (**Fig. 2E**). Next, we specifically measured differences in proportion of each epithelial subset across the four study groups, broadly grouping subsets as basal cell clusters (basal, basal activated, basal cycling), epithelial development clusters (parabasal, early progenitor and club precursor subsets), secretory cells (club and goblet cells) and ciliated development (deuterosomal and ciliated precursor and mature epithelial cells) (**Fig. 2F**). There was a relative decrease in basal cycling cells in both wheeze groups compared to control and no wheeze/RSV groups, and expansion of early progenitor and precursor cells in all groups relative to controls. We also found notable changes in composition of goblet and club epithelial cell clusters that were specific to wheeze groups, while the epithelial cells from the RSV only and control groups were similar (**Fig. 2F**). In summary, ALI culture represented well the differentiation pattern of the epithelium enabling comparative study of epithelial clusters across the four study groups. Together, cluster comparisons across study groups suggested a specific developmental defect common to the wheeze phenotype, while changes related to RSV infection included expansion of early progenitor and precursor cells, but differences compared to control were not as pronounced as in the wheeze sub-groups.

**Fig. 2.**
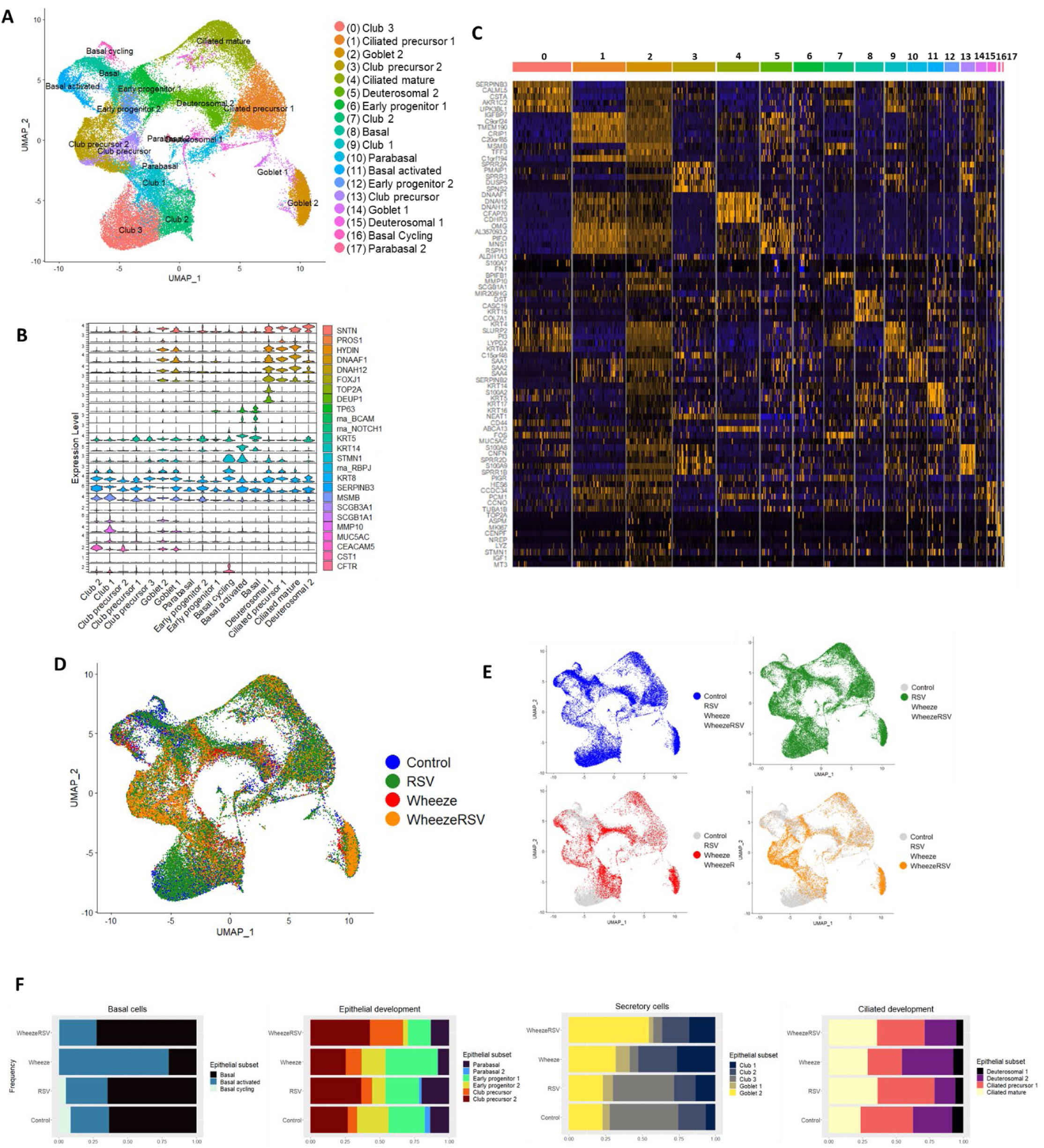
Epithelial subset composition of the developing NAECs (2-3 y.o) in air-liquid interface culture. **A.** An integrated object from all four study groups showing identified epithelial cell subsets. **B.** Marker panel used to identify epithelial cell subsets. **C.** Top markers for each of the seventeen clusters identified by differential gene expression analysis. Cluster numbers on top correspond to cluster numbering in panel (**A**). Low expression shown in blue, high expression in orange. **D.** An integrated object from all four study groups (Wheeze/RSV, Wheeze/No RSV, No Wheeze/RSV, No Wheeze/No RSV [control]) showing cell subset differences by group. **E.** An integrated object from all four study groups highlighting cells individually by group. **F.** Proportional representation of cells per cluster by study group. From left to right: basal cell clusters, progenitor and precursor clusters, secretory clusters and ciliated epithelial clusters. Study groups: Wheeze/RSV in infancy, Wheeze (Wheeze/No RSV), RSV (No Wheeze/RSV in infancy), and Control (no wheeze and no RSV in infancy).

### Characterization of normal versus wheeze airway epithelial phenotypes and differentiation trajectories in air-liquid interface

We next sought to determine whether there are specific differences of epithelium from children with a history of wheeze that distinguish it from healthy (control) epithelium. Our data mining of human birth and early life lung gene expression available from a publicly available resource (LungMAP database) demonstrate that epithelial postnatal development follows a specific trajectory with predominant basal gene expression at birth followed by increasing expression of cell junction and secretory signatures during the first year of life, and signature of ciliation increasing after the first year of life (**Fig. 3A**). The mature epithelial barrier is known to be highly heterogeneous and is comprised of multiple subsets with specialized developmental, structural, secretory, sensory and defense roles in the conducting airways (**Fig. 3B**). Air-liquid interface culture conditions allow for *in vitro* study of a basal-to-specialized epithelial differentiation process, providing the opportunity to compare differentiation trajectories in control vs wheeze AEC phenotypes. Relative proportions of all epithelial subsets across our four study groups suggest a relative increase in all developmentally active cells (basal activated, parabasal, early progenitor, and club cells) in the RSV-only group (no wheeze/RSV) relative to controls but these increases were even greater in wheeze groups. The most dramatic expansion of developing epithelium was observed in epithelial cells derived from children that had both RSV and wheeze in the first year of life (**Fig. 3C**). Cell cycle analysis confirmed progenitor and precursor populations (expressing markers of G2/M and S cell cycle phase) had increased proliferation capacity and these populations were prevalent in children with wheeze (**Fig. 3D**). Slingshot reconstruction of differentiation trajectories of the epithelium revealed significant differences in the development of epithelial cells from the wheeze subgroups relative to control. This suggests that wheezing illnesses induce an alternative precursor brachiation event, and this distinct developmental pathway likely results in alternate mature subsets with functional differences (**Fig. 3E**). In summary, despite common origins in basal cells, airway epithelium from children with wheeze is characterized by differentiation along an altered trajectory with early basal activation of developmental pathways, plasticity in precursor differentiation and the delayed onset of maturation (summarized in **Fig. 3F**).

**Fig. 3.**
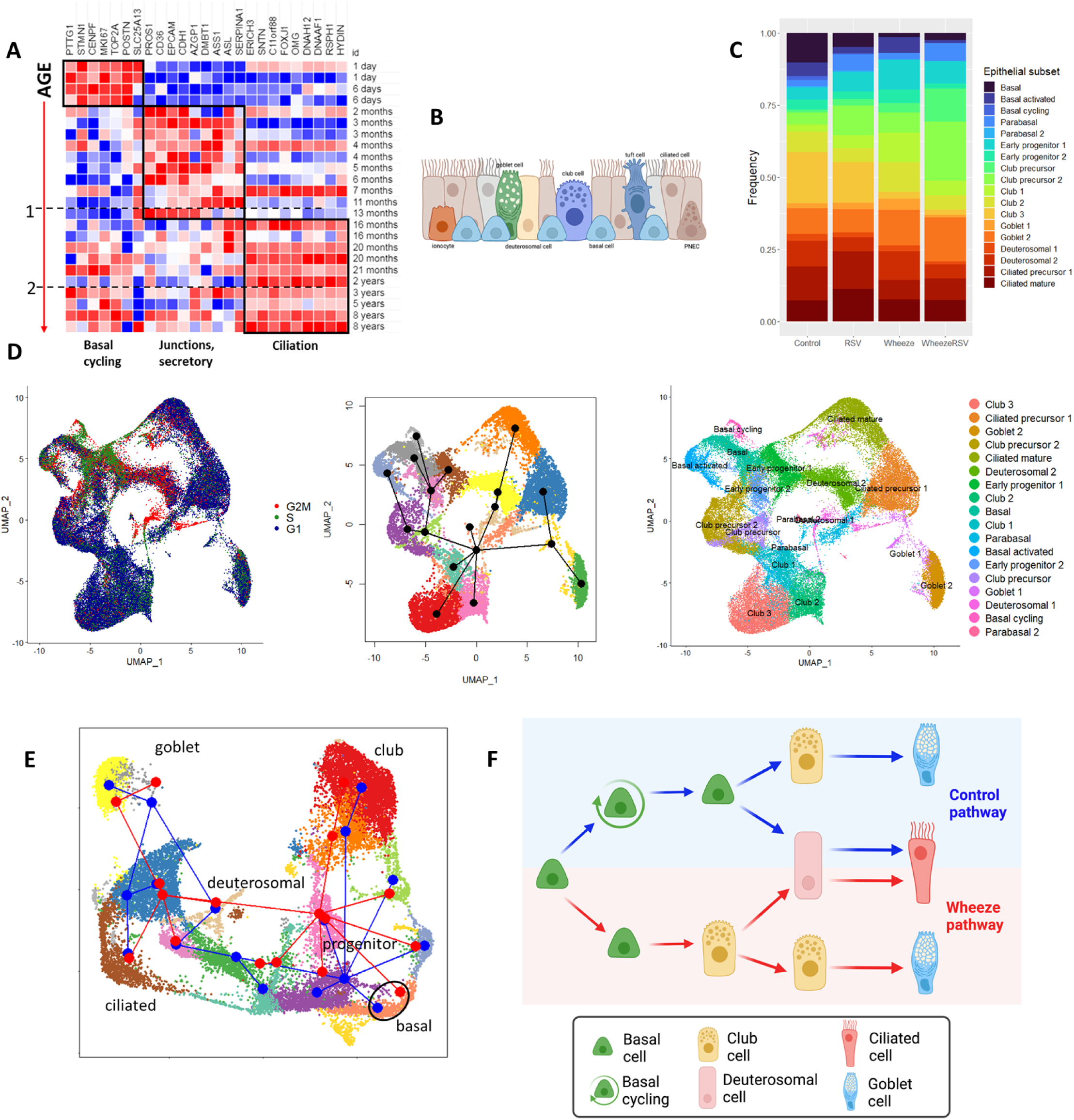
NAECs from wheeze study groups show evidence of developmental reprogramming and an increased expansion of epithelial precursor phenotypes. **A.** LungMAP data mining of human lung tissue development suggests specific epithelial differentiation trajectory in early life. B. A diagram showing heterogeneous composition of mature stratified respiratory epithelium. C. NAECs from wheeze study groups show relative expansion of epithelial progenitor and precursor subsets. D. Cell cycle analysis confirms early identity and two radiation events in basal and progenitor epithelial cell subsets. E. Slingshot developmental trajectory inference reveals altered developmental trajectory in wheezers. F. Interpretation of developmental trajectories in control and wheeze study groups. Panel F created with BioRender.com

### Developmental programming and activation of basal cells in airway epithelium from children with wheeze and RSV infection in the first year of life

Since basal cells represent a starting point in epithelial differentiation, we next examined whether gene expression and biological processes in NAECs collected at age 2-3 years from infants with RSV infection in the first year of life or wheeze differ from controls (No wheeze/No first year RSV infection) specifically in the basal cell subsets. First, we found increased expression of the developmental pathway (WNT, Notch, EGF, TGF, tissue plasminogen) markers specifically in the two wheeze groups (Wheeze/No RSV and Wheeze/RSV) relative to controls (No Wheeze/No RSV), while these markers were not different between RSV only and control groups (**Fig. 4A**). These markers were increased specifically in basal and basal activated subsets, while basal cycling cells were lacking in wheeze groups. For example, expression of PAI-2 (SERPINB2) and JAG1 (Notch/Jagged) was increased specifically in wheeze NAECs (**Fig. 4B**).

**Fig. 4.**
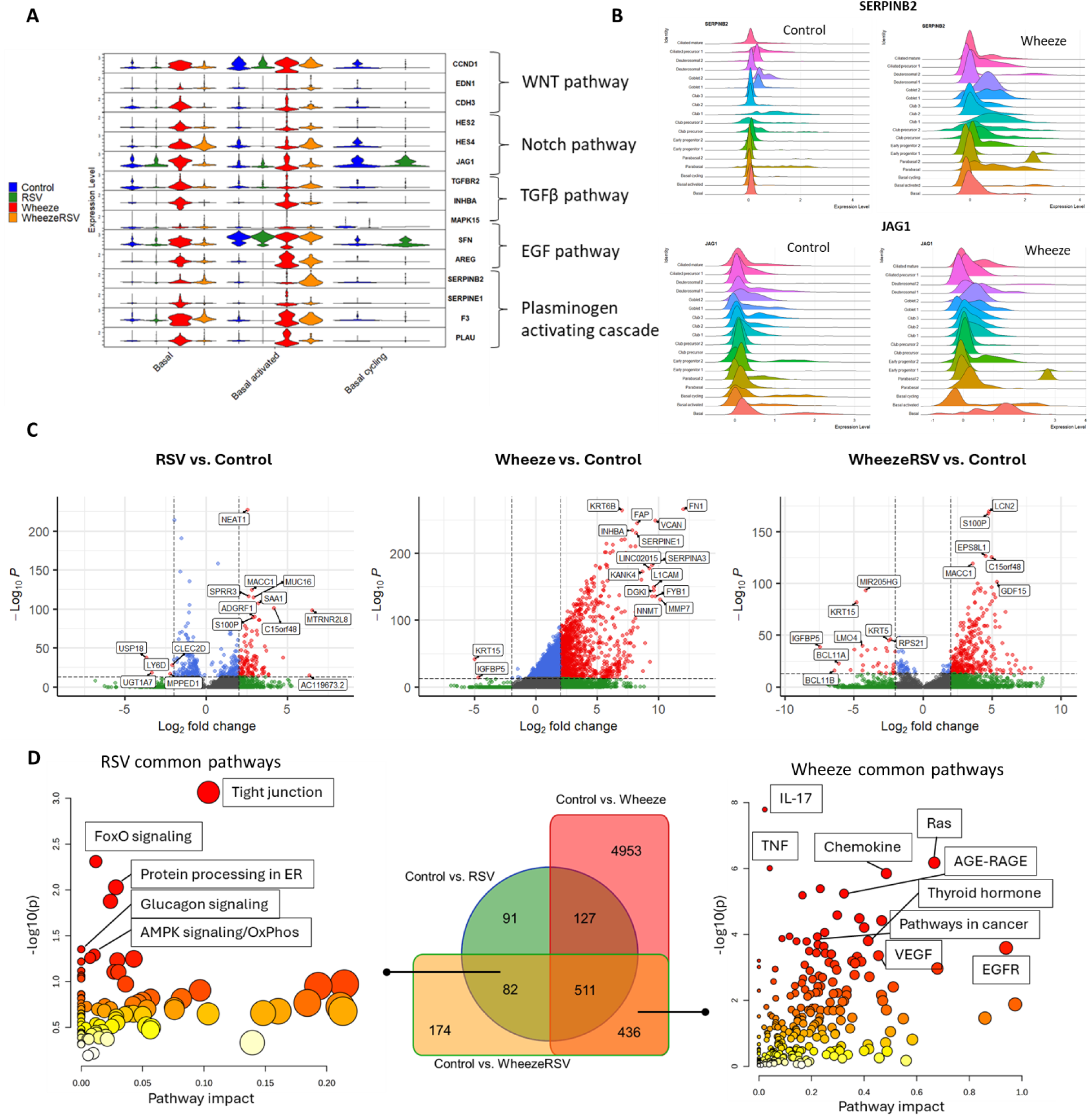
NAECs from wheeze study groups show early activity of developmental pathways and abnormal activation of basal cells. **A.** Expression of markers of developmental pathways (WNT, Notch, TGF, EGF, tissue plasminogen system) by cell type and study group. **B.** Expression of PAI-2 (SERPINB2) and JAG1 (Notch/Jagged) is increased in wheeze NAECs. **C.** Differentially expressed gene (DEG) analysis of basal cell subset comparing RSV and wheeze study groups to controls. FDR-adjusted p-values used. **D.** Pathway analysis of upregulated basal cell DEGs common to “Control vs. RSV” and “Control vs. WheezeRSV” (left) and upregulated basal cell DEGs common to “Control vs. Wheeze” and “Control vs. WheezeRSV” (right).

Next, we determined differentially expressed genes (DEGs) between each of our study groups relative to the control group (**Fig. 4C**). Wheeze and Wheeze/RSV groups had the highest number of upregulated genes relative to controls, while the No wheeze/RSV group had the lowest number of DEGs compared to controls (No wheeze/No RSV). Top upregulated genes in NAECs from infants with first year RSV infection included Nuclear Enriched Abundant Transcript 1 (NEAT1), MET transcriptional regulator MACC1, mucin MUC16 and small proline rich protein 3 (SPRR3). Top genes in NAECs from children with wheeze included fibronectin (FN1) and keratin 6B (KRT6B), while the Wheeze/RSV group highly expressed lipocalin-2 (LCN2) and S100 calcium binding protein P (S100P). Interestingly, keratin KRT15 and IGF binding protein 5 (IGFBP5) were commonly downregulated in wheeze groups with and without infant RSV infection (**Fig. 4C**).

Next, to understand specific differences between wheeze and RSV basal cell activation, we performed a pathway analysis on upregulated basal cell DEGs common to “Control vs. RSV” and “Control vs. Wheeze/RSV” comparisons (thus representing RSV-only response) and upregulated basal cell DEGs common to “Control vs. Wheeze” and “Control vs. Wheeze/RSV” comparisons (representing wheeze-specific biology) (**Fig. 4D**). We found that RSV basal activation is associated with pathways that include tight junction regulation, FoxO signaling pathway, protein processing in endoplasmic reticulum, glucagon signaling, AMPK signaling and oxidative phosphorylation. Wheeze phenotype is characterized by basal cell activation and upregulation of immune signaling pathways (IL-17, TNF, chemokines), RAS AGE-RAGE, thyroid signaling, and growth factor/remodeling pathways (VEGF, EGF, pathways in cancer (WNT, Notch/Jagged)), among others.

Together, these findings show that the wheeze phenotype is associated with significant immune activation, remodeling and abnormal developmental activity of basal cells, which likely sets wheeze NAECs on an aberrant differentiation trajectory, whereas infant RSV infection alone is associated with a more subtle impact on epithelial cellular processes that would lead to disruption of barrier development.

### Biological processes associated with extracellular matrix deposition and immune activation in NAECs by wheeze phenotype

Next, we performed an enrichment analysis to determine which biological processes and transcriptional programs are associated with aberrant development of NAECs from children with wheeze. **Fig. S2A** shows biological enrichment analysis of genes expressed in different developmental clusters comparing control and wheeze only-groups. This analysis showed increased gene activity associated with remodeling of the extracellular matrix in basal cells from the wheeze group consistent with the relative increase in basal activated cells noted above. Notably, we also found evidence for sustained immune activation of epithelial cells from children with wheeze, specifically in parabasal, early progenitor and goblet cells, evidenced by expression of chemokines, cytokines and antigen presentation genes. These processes were not prominent in the NAECs from control children, which only showed processes consistent with normal epithelial development, keratinization and junction formation (**Fig. S2A**). Transcriptional factor inference analysis confirmed these functional differences suggesting early activation of BRCA1, Myc, TGFb pathway (SMADs) in children with wheeze and sustained progenitor transcription programs (JUN, JUND/B) (**Fig. S2B**). We specifically noted increased expression of panel of gene markers associated with remodeling (SERPINB2, THBS1, TGFBI, TNC, COL1A1, VCAN, FN1, SPP1, TAGLN, POSTN) in airway epithelium of children with wheeze (**Fig. S2C**).

Notably, NAECs from children with wheeze expressed KRT13 (marker of hillock cells) in early progenitor and club precursor subsets along with CXCL8 (immune) and CFB (complement) markers, while remodeling markers such as FN1 (fibronectin) noted above were expressed in basal and early progenitor subsets (**Fig. S2D**). In summary, these results show that the wheeze epithelial developmental phenotype is characterized by aberrant activation of basal and club precursor (hillock) cells, and persistent activation of remodeling, complement and immune pathways during development.

### Diversity of RSV receptors, expression of anti-viral response genes, barrier permeability and susceptibility to *in vitro* RSV infection in NAECs from wheeze and no wheeze sub-groups

Intriguingly, NAECs from wheeze groups showed increased diversity of all currently known RSV receptors, which were increased across different epithelial subsets in RSV and wheeze subgroups (**Fig. 5A and B**). RSV receptor genes for this panel were determined based on previously published reports (*19, 20*). Moreover, NAECs from wheeze groups had decreased expression of MX1 (MX Dynamin Like GTPase 1) and IFNAR1 (Interferon-alpha/beta receptor alpha chain), key markers of cellular antiviral responses (**Fig. 5C and D**).

**Fig. 5.**
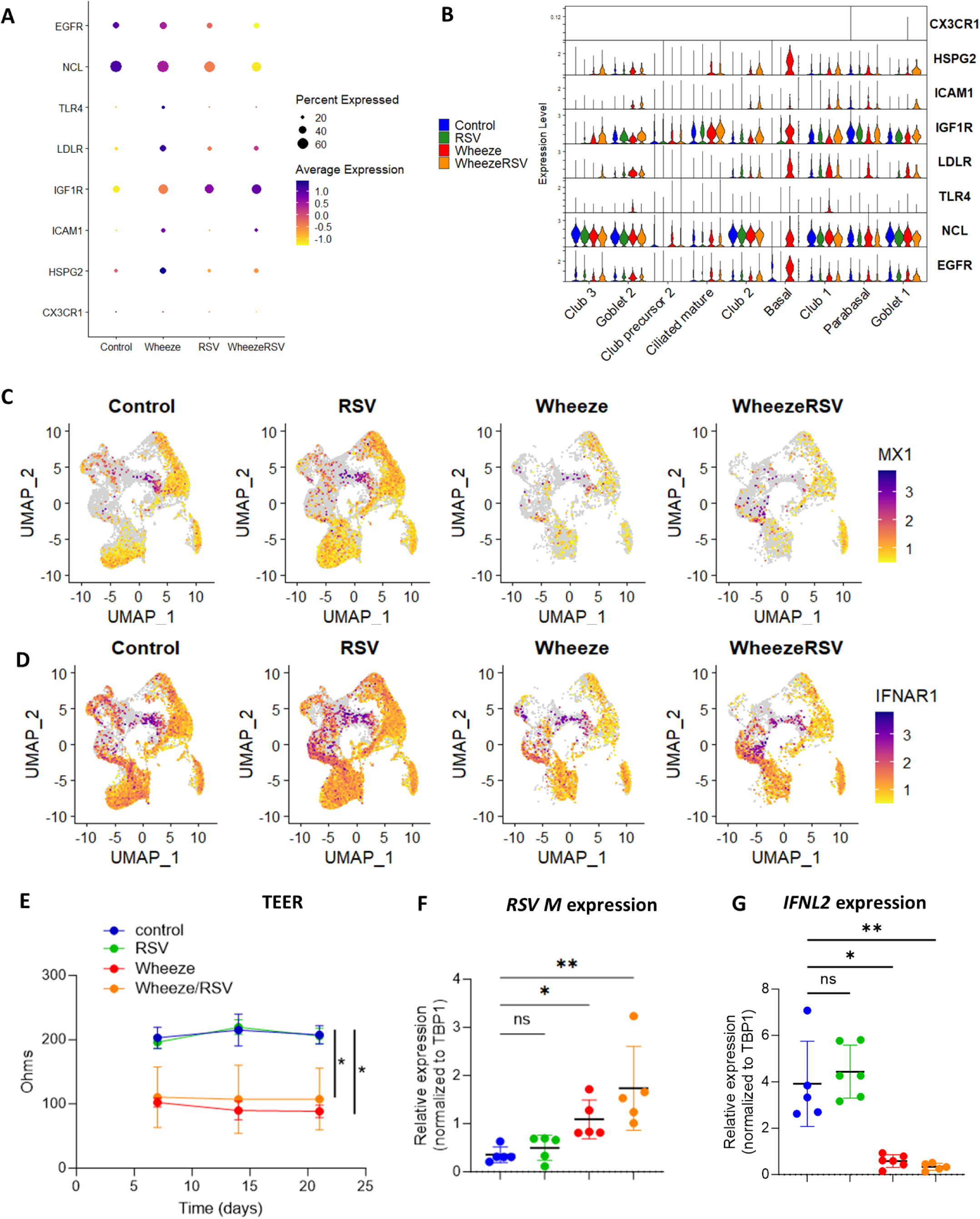
NAECs from wheeze study groups show increased diversity of RSV receptors, decreased expression of host anti-viral response genes, increased barrier permeability and increased susceptibility to RSV infection in vitro. **A and B.** NAECs from wheeze groups show increased diversity of RSV receptors. **C.** NAECs from wheeze groups have decreased expression of MX1 (MX Dynamin Like GTPase 1), a marker of cellular antiviral response. **D.** NAECs from wheeze groups have decreased expression of IFNAR1 (Interferon-alpha/beta receptor alpha chain), another antiviral factor. **E.** NAECs were differentiated at air-liquid interface TEER permeability was determined at various days post air-liquid interface. n=5-6 in each group and data is shown as mean (SD), * p<0.05, two-way ANOVA with repeated measures with Tukey post-hoc test. **F-G.** After fully differentiated, NAECs were infected with RSV 01/2-20 (MOI=3). Twenty-four hours post infection, cells were harvested and RSV M gene expression (panel D) or IFNL2 (panel E) was determined by qPCR and normalized to TBP1. n=5-6 in each group and data is shown as mean (SD)* p<0.05 ** p<0.01, one-way ANOVA with Tukey post-hoc test; ns, not significant.

Based on these data, we wanted to explore if differentiated NAECs had different RSV infectivity in culture. To do this, NAECs from 5-6 unique children in each of the 4 *a priori* selected groups (No Wheeze/No RSV [control], Wheeze/No RSV, Wheeze/RSV, and No Wheeze/RSV) were differentiated. Starting at the time of air-liquid interface (day 7), TEER permeability was assessed as a measure of barrier capacity of the NAEC cultures from the 4 groups. TEER permeability measurements were reduced in NAECs from wheeze and wheeze/RSV children compared to NAECs from control or No wheeze/RSV children (**Fig. 5E**). At day 21 of ALI, after full differentiation, NAECs were infected *in vitro* with a clinically relevant strain of RSV (RSV 01/2-20) at a MOI=3 or mock infection. Expression of RSV M gene was determined 24 hours post infection, and NAECs from wheeze and wheeze/RSV children had increased RSV M gene expression compared to NAECs from control and no wheeze/RSV children (**Fig. 5F**). IFN production is known to increase after RSV infection as an initial anti-viral response. Therefore, we also measured IFNL2 gene expression. As expected, *in vitro* RSV 01-2/20 infection increased IFNL2 expression compared to mock infected cells for each group (data not shown).

However, NAECs from control (No wheeze/No infant RSV infection) and RSV (No wheeze/RSV) children had significantly increased IFNL expression after *in vitro* RSV 01-2/20 infection compared to NAECs from children with wheeze/No RSV infection and wheeze/RSV phenotypes (**Fig. 5G**). Together, these findings suggest that wheeze is an underlying phenotype with potentially genetic or epigenetic origins and is characterized by increased barrier permeability and diversity of RSV receptors, which may predispose to infection, enhanced RSV infection severity in infancy or long-term effects of RSV infection.

## Discussion

Using nasal airway epithelial cells from children who had careful characterization of RSV infection and wheeze in the first year of life and single cell profiling of these cells collected at the end point of postnatal differentiation (age 2-3 years), we demonstrated that nasal airway epithelial cells from children with a wheeze phenotype are characterized by an altered air-liquid interface differentiation trajectory in culture with early basal activation of developmental pathways, plasticity in precursor differentiation and persistent expansion of developmentally active epithelial cell subsets. NAECs from children with wheeze also have increased diversity of all currently known RSV receptors and blunted anti-viral immune responses to *in vitro* infection. Together our findings suggest that airway epithelial developmental reprogramming in children with wheeze has a genetic or epigenetic origin which is characterized by increased barrier permeability and altered antiviral response in addition to increased diversity of RSV receptors which may predispose to and amplify the effects of RSV infection in infancy. Early life RSV infection results in an additive effect on aberrant airway epithelial development most pronounced in those with a wheeze phenotype, suggesting a potential gene by environment interaction that may contribute to and explain the association of early life RSV infection with childhood asthma risk.

One of the remarkable findings in our study was the level of epithelial developmental reprogramming associated with an early life wheeze phenotype compared with non-wheezers. Airway epithelial cells from children with an early life wheeze phenotype (wheeze with or without RSV) also had more pronounced epithelial reprogramming than cells from children that had RSV in infancy without wheeze. Our findings point to developmental defects originating in basal cells of children with wheeze, which include increased expression of developmental pathways (WNT, Notch/Jagged, TGFβ, EGF), tissue plasminogen system activation and increased production of extracellular matrix proteins (versican, fibronectin, periostin, tenascin C, among others). Interestingly, markers such as periostin and PAI-2 also traditionally serve as markers of “Type 2-high” molecular endotype of allergic asthma. The early appearance and persistence of developmental defects in these pathways in epithelial culture among children with a wheeze phenotype may suggest developmental/epigenetic origins of asthma that might initiate persistent airway inflammation in some children and promote the development of allergic inflammation. In support of this concept, we and others have reported dysregulation of epithelial developmental pathways in wheeze and adult asthma (*21–24*).

Nasal airway epithelial culture in air liquid interface allowed for the *in vitro* assessment of epithelial developmental processes from basal to specialized differentiated cells. If terminology of heterochrony (study of differences in the timing, rate, or duration of a developmental process) were to be used, the effect of basal reprogramming on the wheeze NAEC differentiation could be characterized as an “early developmental onset – delayed developmental offset” hypermorphosis (“hyperdevelopmental” trajectory)(*25*). Our lineage reconstruction analysis demonstrated persistence and proliferation of specific epithelial cell subsets including early progenitor and club precursor epithelial cells. Our investigation of club precursor cells from children with a wheeze phenotype determined that they express KRT13, which in conjunction with club precursor markers would classify them as recently discovered “hillock” cells (*26*). Little is currently known about the role of hillock cells in wheezing illnesses and asthma. Montoro et al. described them as a variety of club cells descending from the basal cell lineage and expressing keratin 13. These cells are found in contiguous groups of stratified epithelial cells, forming structures termed ‘hillocks’, where they have high cycling capacity and express markers associated with squamous epithelial differentiation, cellular adhesion and immunomodulation (*26, 27*). These cells were also recently reported in a human model of asthma exacerbation *in vitro* associated with an allergic asthma phenotype (*28*). Since this transient population between basal and club cells was associated with abnormal epithelial differentiation, we can deduce that this reprogrammed epithelial subset is playing a role in perpetual epithelial barrier injury-repair/remodeling processes (or a transient role in normal wound healing). Our findings suggest that hillocks are part of a broader developmental reprogramming following preschool wheezing illnesses and could be markers of abnormal barrier development in early life. Moreover, abnormal wheeze differentiation was associated with persistence of complement system and immune gene expression (i.e., IL32, CXCL3, CXCL8, HLA-DRA, HLA-DRB1, HLA-DQA1, CFB) in different epithelial subsets (parabasal, early progenitor, goblet). This may indicate a baseline inflammatory state of the epithelium even in the absence of active interaction with immune cells (*29, 30*) and may lead to abnormal immune responses at the epithelial barrier interface.

The impact of RSV on *in vivo* differentiation of epithelium early in life and the development of asthma had been unexplored. RSV can reprogram differentiation of AECs i*n vitro* by infecting and altering the developmental course of epithelial progenitor cells (*31*). However, our study uncovered the significance of a pre-wheeze epithelial developmental phenotype as a potential predisposition factor for aberrant barrier function, enhanced risk for RSV infection, and perhaps infection severity. While we describe broader changes in cell types and pathways associated with the wheeze phenotype (basal subset activation, mucociliary precursor expansion) than with infant RSV infection, in combination there appears to be an additive effect on aberrant airway epithelial development. It is critical to determine which factors drive this early susceptibility.

Interestingly, we also measured greater expression of RSV receptors on epithelial cells of children with an early wheeze phenotype. Some of the suspected RSV-associated receptors (IGF1R, heparan sulfate proteoglycan, EGFR) are also active participants in epithelial and matrix remodeling, which suggests that barrier remodeling processes that we identified as part of the wheeze phenotype may drive increased receptor expression for RSV. Moreover, we found downregulation of anti-viral response markers, such as MX1 and IFNAR1, in the epithelial cells from children with wheeze, which suggest that developmental reprogramming in wheeze impacts the ability of the airway epithelium to mount anti-viral defense. This finding in epithelium is consistent with our finding of suppressed anti-viral immunity in PBMCs from children with prior RSV infection in infancy (*32*) and previous reports in childhood allergic asthma (*33*). The functional consequences of this are supported by our findings of increased *in vitro* RSV infection as measured by RSV *M* gene expression and decreased expression of *IFNL2* in NAECs from children with an early life wheeze phenotype compared to NAECs from non-wheezers. Further, children with both an early life wheeze phenotype and infant RSV infection had the most dramatic changes in epithelial differentiation and function relative to controls. The distinct differences in pathways associated with the wheeze phenotype and early life RSV infection may suggest a “double hit” scenario in epithelial development in infancy: an early wheeze phenotype at or after birth due to genetic or epigenetic modification with increased susceptibility to RSV, and early life infection further amplifying developmental defects in early life and leading to differentiation to a functionally impaired airway barrier. This also supports a shared genetic predisposition for both wheeze/asthma and more severe RSV infection which we have previously demonstrated (*34*). An airway developmental trajectory characterized by barrier disruption and enhanced susceptibility to infection may predispose a child to increased immune mucosal surveillance, higher penetrance and processing of environmental antigens, leading to atopy and allergic disease, as well as enhanced susceptibility to asthma risk factors such as secondhand smoke and air pollution.

The novelty of this study is the use of primary cells collected from children during well-child visits at age 2-3, representing the end point of postnatal epithelial differentiation, who had careful characterization of both RSV infection and wheeze in early life. Limitations of the study include cross-sectional collection of airway epithelial cells for the main analysis. We performed scRNA-seq analysis of 2-3 year-old NAEC differentiation in ALI culture, which captured the basal-to-mature re-differentiation process rather than a childhood epithelial developmental curve or *in situ* differences between established airway subsets. Small sample size is another limitation, which limited power to observe differentially expressed genes. While epithelial cell populations functionally differ between the upper and lower airway, basal-to-specialized epithelial differentiation pathways are remarkably similar in terms of developmental processes in the upper and lower airway, and transcriptomic data support the important parallels between the upper and lower airway epithelium. This supports the use of NAECs in children to model the differences in airway epithelial development that characterize wheeze and respiratory viral infection (*13, 14, 35*).

This study provides new evidence for how the airway epithelium develops in disease, and how and why it responds differently to environmental stress. Understanding development in the airway epithelial barrier after birth through early childhood is recognized as a key to understanding the developmental origins of childhood wheeze and asthma. The airway epithelium is a fundamental mucosal barrier, and the nasal epithelium provides critical first responses against viral infections and illnesses. The findings of our study suggest that RSV infection and wheezing illnesses alter nasal airway epithelial development including barrier function and the response to respiratory viruses. We postulate that wheezing illnesses and early life RSV infection might change in the airway epithelium in some children to enhance susceptibility to respiratory viral infections thus increasing the risk of additional wheezing illnesses and possible chronic airway obstruction. As there are no effective primary preventive interventions for asthma, identifying the timing and pathways driving airway epithelial development may inform novel targets for prevention and treatment approaches that regulate the normal development of the early life airway epithelium with the potential to prevent both severe respiratory viral infections and asthma.

## Materials and methods

### Study population

The Infant Susceptibility to Pulmonary Infections and Asthma Following RSV Exposure study (INSPIRE) is a large, population-based birth cohort of healthy, term children (n=1,946) specifically designed to test the association of RSV infection in infancy with risk of childhood asthma. RSV infection status (uninfected *vs.* infected) was ascertained in the first year of life using a combination of passive and active surveillance with viral identification through molecular and serological testing. Children were followed prospectively for annual recurrent wheeze using a validated questionnaire (*16, 17, 36*). The study population in this proposal is an *a priori* designed nested cohort of 100 participants selected for follow-up using a random number generator from 4 groups of children with and without wheezing, and RSV infected and uninfected during infancy (no wheeze/no RSV [controls], no wheeze/RSV, wheeze/RSV, wheeze/no RSV) (**Fig. S1**). These children completed additional in-person study visits between age 2-3 years that included NAEC collection and culture representing the end point of postnatal airway epithelial differentiation. From this nested cohort of 100 children, two to three participants in each of these 4 groups were randomly selected. We selected samples from children with wheeze who had wheeze during follow-up between ages 1-4. All NAECs were differentiated in ALI culture and sent for scRNA-seq. In additional experiments, five to six individual participants were selected from each of the 4 groups and NAECs were differentiated and TEER was measured during differentiation. These NAECs were infected with RSV 01/2-20 or mock for follow-up studies. The Institutional Review Board of Vanderbilt University Medical Center approved this study and one parent of each child provided informed consent for their participation. The detailed methods for INSPIRE have been previously reported (*36*).

### Human nasal airway epithelial cell culture

NAECs were collected from the *a priori* nested cohort of INPSIRE children during a well-child visit between 2-3 years of age. Children were screened for signs and symptoms of respiratory illness, and if detected, visits and collections were rescheduled. NAECs were collected from subjects by brushing nasal passages at the level of the inferior turbinate with a soft flocked cotton swab (Copan, Murrieta, CA, USA) and placing them in cold PneumaCult-Ex Plus Medium (StemCell, Vancouver, Canada). Cells were placed on collagen coated flasks and then were submerged and expanded in PneumaCult-Ex Plus Medium, consisting of 500 mL Ex Plus Basal Medium supplemented with 10 mL PneumaCult-Ex Plus 50X supplement, 0.5 mL Hydrocortisone stock solution, and Pen-Strep. Once cells reached 50-70% confluence on the flask, NAECs were disassociated using 2-4 mL Animal Component-Free (ACF) Cell Dissociation Solution and 2-4 ml ACF Inhibition Solution (StemCell, Vancouver, Canada) and transferred to a 24 transwell plate with transwells coated in 200 µL of rat tail collagen I. Cells remained submerged in PneumaCult-Ex Plus Medium until confluency (approximately 1 week) with media changed every other day. Once confluent, media was removed from the apical chamber and the NAECs were allowed to differentiate for 3 weeks using PneumaCult ALI (air-liquid interface) Medium (500 uL) in the basal chamber. Media in the basal chamber was changed every other day. Complete PneumaCult ALI Medium consists of 490 mL ALI Medium plus 50 mL of PneumaCult-ALI 10X supplement, 5 mL of PneumaCult ALI maintenance supplements, 2.5 mL Hydrocortisone stock solution, and 1% Pen-Strep. Laboratory staff involved in NAEC culture, TEER measurements and *in vitro* RSV infection were blinded to group assignment.

### Single cell RNA-Seq

For each sample sequenced (N=2-3/group), 10,000 cell target was used for single-cell capturing and library construction using Chromium Single Cell 3’ Reagent Kits (v3.1 chemistry, PN-1000130) from 10x Genomics, according to manufacturer’s instructions. Single-cell Gel Beads-in-Emulsion (GEMs) captured cells underwent lysis and transcript barcoding. The corresponding cDNA along with cell barcodes were amplified using PCR. scRNA-seq libraries were constructed using 10x Genomics Library Construction kits (PN-1000196) and Dual Index Kit TT Set A (PN-1000215). The constructed libraries were sequenced on an Illumina platform to generate paired end reads. The resulting raw sequencing data was processed using the CellRanger pipeline (version 7.1.0, 10x genomics).

### In vitro RSV infection of NAECs

After NAECs were fully differentiated (>21 days at ALI), each of the donor cells were infected on the apical surface (top chamber) with 50 µL of RSV 01/2-20 (MOI = 3) or mock. After infection, cells were incubated for an hour at 37°C with gentle rocking. After an hour, inoculum was removed from the apical surface, cells were washed 1X with PBS, and cells were placed back into the appropriate temperature incubator. Basolateral supernatant, viral washings, and cells were collected at 24 hours post infection.

### TEER measurements

The barrier function of NAECs was determined by measuring the trans-epithelial electrical resistance (TEER) as previously described (*37*) at days 7, 14, and 21 of ALI differentiation. Cells were allowed to equilibrate to room temperature for 15 minutes prior to TEER measurement. ALI media was replaced by PBS on the basolateral side and PBS was added to the apical side of each transwell prior to TEER measurement. TEER was measured under a sterile hood with a Chopstick electrode and an epithelial voltammeter (EVOM^2^) (World Precision Instruments, Sarasota, FL). Each well was measured in triplicate and values were averaged. After measurement, PBS was removed, and ALI media was replenished on the basolateral side.

### RNA isolation, cDNA generation, and quantitative PCR (qPCR)

For RNA isolation, lysed cells were thawed then passed through a QIAshredder (Qiagen, Hilden, Germany). RNA was then extracted using the RNeasy Mini Kit (Qiagen, Hilden, Germany). RNA quality was assessed using a NanoDrop 2000 Spectrophotometer (Thermo Fisher Scientific, Waltham, USA). cDNA was generated using the SuperScript IV First-Strand Synthesis kit (Thermo Fisher Scientific, Waltham, USA). qPCR was conducted using QuantStudio and gene expression was normalized to a housekeeping gene of TBP1. RNA M gene primers (forward: GGC AAA TAT GGA AAC ATA CGT GAA; reverse: TCT TTT TCT AGG ACA TTG TAY TGA ACA) and were generated using IDT. IFNL2 gene primer (catalog number: 4331182; assay ID: Hs00820125_g1) and Housekeeping gene TBP primer (catalog number: 4331182; assay ID: Hs00427620) were purchased from Thermo Fischer Scientific (Waltham, MA, USA).

### Bioinformatic and statistical analysis

Seurat R package version 4.1.1 was utilized for analyzing counts data, normalization, dimension reduction, clustering, integration, visualization, identification of unique cluster markers and differential gene expression analysis. Cell cycle analysis was performed utilizing *cc.genes* function in Seurat. *Slingshot* v.2.2.1.R package was used for differentiation trajectory inference. *Ggplot2* R package version 3.4.0 was used for figure generation and visualization. Differentially expressed genes (DEG) were determined using non-parametric Wilcoxon rank sum test implemented in Seurat. *EnhancedVolcano* R package was used to generate volcano plots in **Fig. 4C** based on Seurat DEG analysis. The data mining results shown in **Fig. 3A** are based upon human lung RNA-seq data (neonate, infant and child) generated by the LungMAP Consortium and downloaded from (www.lungmap.net), in 2020. The LungMAP consortium and the LungMAP Data Coordinating Center (U24-HL148865) are funded by the National Heart, Lung, and Blood Institute (NHLBI) (*38*). Pathway analysis in **Fig. 4D** was performed in Metaboanalyst (*39*) (“genes only” option) using KEGG pathway reference and hypergeometric test-based enrichment analysis. We obtained biological processes in **Fig. S2A** associated with the significant genes identified from our differential expression analysis using Enrichr (*40*) with Reactome and KEGG based pathway analysis with hypergeometric statistical testing and *Clustergrammer* hierarchical clustering. TEER in **Fig. 5E** was analyzed using two-way ANOVA with repeated measures and a Tukey post hoc test and presented as a line graph depicting the mean ± the standard deviation (SD) with colors of groups shown in the legend. The qPCR data in **Fig. 5F and G** was analyzed using one-way ANOVA with a Tukey post hoc test and presented as bar graphs depicting the mean ± the standard deviation (SD) with sample number listed in each figure legend.

### Data and code sharing statement

Sequencing data will be made available in GEO NCBI and ArrrayExpress (EMBL-EBI) databases as well as upon request. R code used to analyze data will be shared upon request.

## Acknowledgements

We are deeply grateful to all of the families who participated in this study, and to the middle Tennessee pediatric practices with whom we collaborated to enroll a representative population of our region. We are also appreciative of Christine Cole Johnson, PhD and Alexandra Sitarik, PhD for their critical review of the manuscript.

## Authors’ contributions

SB, DN, LA and TH contributed to the conception of the study and/or the study design. TH, SB, JG, LA and DN obtained the research funding supporting this study. TH and TG designed and conducted the clinical study and TH oversaw biospecimen acquisition. TH, TG and CR oversaw primary data and validation of clinical outcomes. DN, JC, KM and SK led the nasal airway epithelial culture, cryopreservation, growth in ALI, *in vitro* infection. NH provided technical support. JC contributed to the biospecimen processing, cell culture and laboratory assays. Data processing and/or analyses were performed by SB, CM, SM, and TG. The manuscript and figures were drafted by TH, DN and SB. Co-first authorship was determined based on the equal contributions by SB and DN to the manuscript. SB, DN and TH drafted the manuscript and all authors contributed to critical revision of the manuscript for important intellectual content. All authors read, critically edited the manuscript and approved the final manuscript.

---

**Fig S1.**
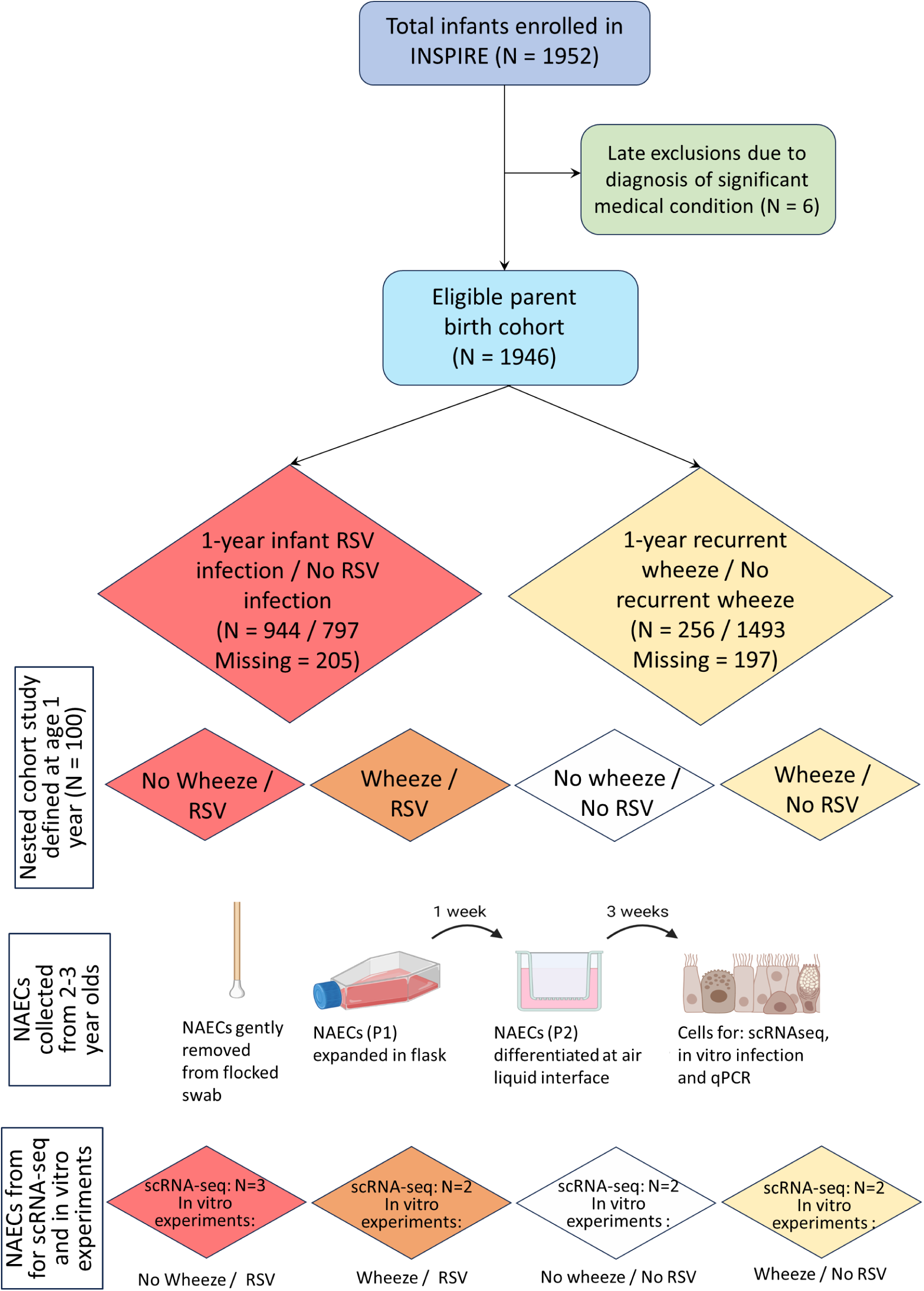
Flow diagram of a priori designed nested cohort study of 100 INSPIRE participants selected into one of four mutually exclusive groups with recurrent wheeze/no recurrent wheeze during the first year of life using validated questionnaires, and RSV infection/no RSV infection during the first year of life as determined by active surveillance using PCR and 1-year RSV serology. These 100 children were seen for an additional research visit at age 2.5 years with collection of nasal airway epithelial cells (NAEC) for culture and RNA-sequencing. A randomly selected group of children with NAEC in each of the four groups representing males and females were randomly selected for single cell RNA-sequencing, and for culture in air liquid interface for in vitro infection with clinical isolates of RSV to measure RSV infectivity and barrier permeability using transepithelial electrical resistance (TEER).

**Fig. S2.**
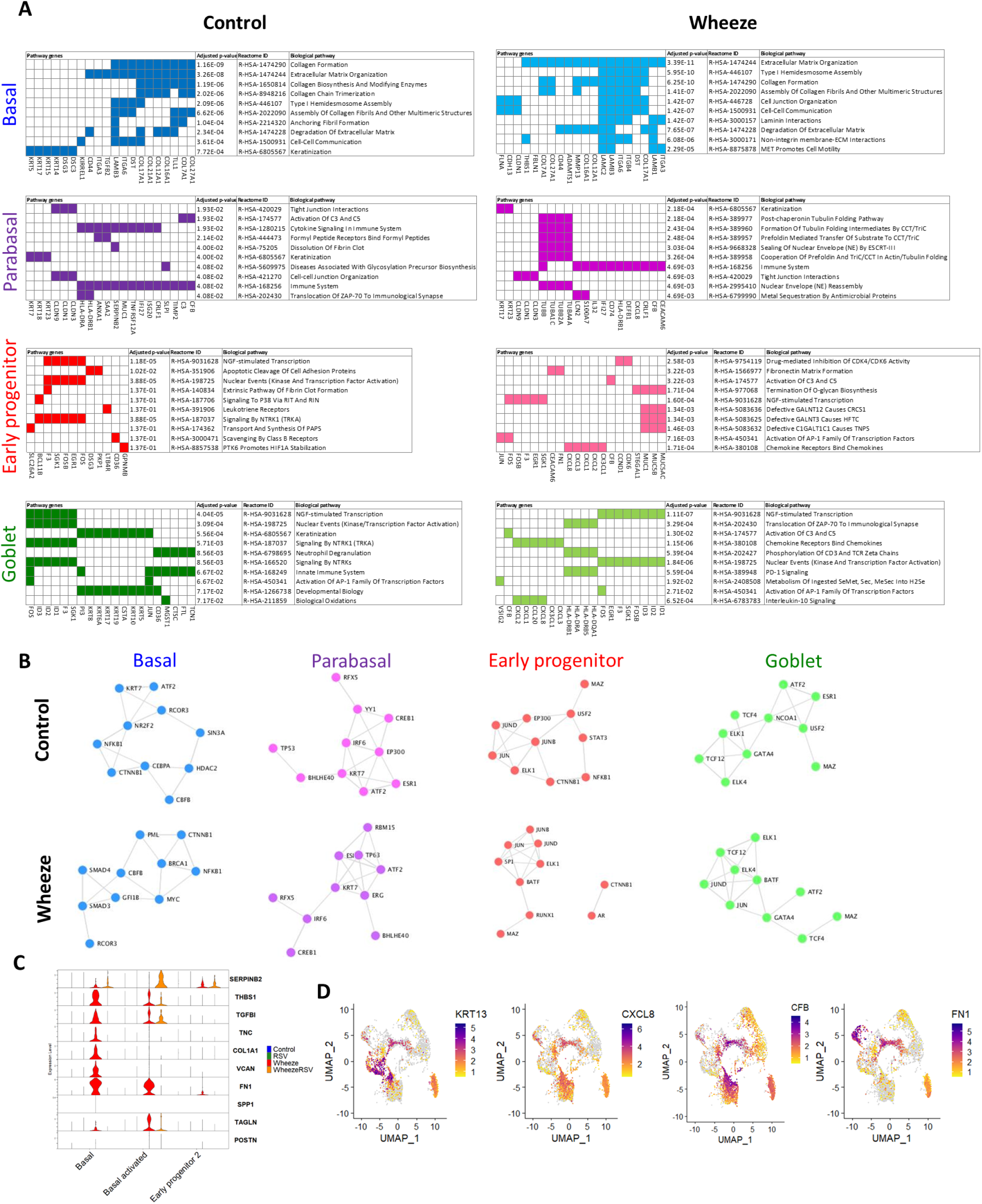
NAECs from wheeze study groups show abnormal and sustained biological processes associated with extracellular matrix deposition and immune activation. **A.** Biological enrichment analysis of genes expressed in different developmental clusters by study group. **B.** Transcriptional factor inference. **C.** Expression of gene markers associated with remodeling, immune and tissue plasminogen system activation in control and wheeze groups. **D.** Expression of KRT13 (hillock cell marker), CXCL8 (immune marker), CFB (complement) and FN1 (fibronectin) in wheeze epithelial cells.

